# LOTVS: a global collection of permanent vegetation plots

**DOI:** 10.1101/2021.09.29.462383

**Authors:** Marta Gaia Sperandii, Francesco de Bello, Enrique Valencia, Lars Götzenberger, Manuele Bazzichetto, Thomas Galland, Anna E-Vojtkó, Luisa Conti, Peter B. Adler, Hannah Buckley, Jiří Danihelka, Nicola J. Day, Jürgen Dengler, David J. Eldridge, Marc Estiarte, Ricardo García-González, Eric Garnier, Daniel Gómez-García, Lauren Hallett, Susan Harrison, Tomas Herben, Ricardo Ibáñez, Anke Jentsch, Norbert Juergens, Miklós Kertész, Duncan M. Kimuyu, Katja Klumpp, Mike Le Duc, Frédérique Louault, Rob H. Marrs, Gábor Ónodi, Robin J. Pakeman, Meelis Pärtel, Begoña Peco, Josep Peñuelas, Marta Rueda, Wolfgang Schmidt, Ute Schmiedel, Martin Schuetz, Hana Skalova, Petr Šmilauer, Marie Šmilauerová, Christian Smit, Ming-Hua Song, Martin Stock, James Val, Vigdis Vandvik, Karsten Wesche, Susan K. Wiser, Ben A. Woodcock, Truman P. Young, Fei-Hai Yu, Martin Zobel, Jan Lepš

## Abstract

Analysing temporal patterns in plant communities is extremely important to quantify the extent and the consequences of ecological changes, especially considering the current biodiversity crisis. Long-term data collected through the regular sampling of permanent plots represent the most accurate resource to study ecological succession, analyse the stability of a community over time and understand the mechanisms driving vegetation change. We hereby present the LOng-Term Vegetation Sampling (LOTVS) initiative, a global collection of vegetation time-series derived from the regular monitoring of vascular plants in permanent plots. With 79 datasets from five continents and 7789 vegetation time-series monitored for at least six years and mostly on an annual basis, LOTVS possibly represents the largest collection of temporally fine-grained vegetation time-series derived from permanent plots and made accessible to the research community. As such, it has an outstanding potential to support innovative research in the fields of vegetation science, plant ecology and temporal ecology.

## 1. Background

Anthropogenic changes are severely impacting our ecosystems (Bradshaw et al. 2021). The rate of species loss has now exceeded background extinction rates (Pimm et al. 2014), leading many scientists to claim a sixth mass extinction (Pereira, Navarro, & Martins, 2012; Ceballos et al. 2015). At the same time, a considerable proportion of natural habitats has been lost (Convention on Biological Diversity 2020) and a number of ecosystem functions and services are seriously at risk (IPBES 2019).

The analysis of time-based patterns in biological communities, especially when focused on primary producers like plants, represents an opportunity to quantify the extent and the consequences of such changes in biodiversity (Dornelas et al. 2014; Gonzalez et al. 2016; Blowes et al. 2019). This research field has potential for: i) unravelling the mechanisms that drive and maintain biodiversity over time (Jones et al. 2017; Hillebrand et al. 2018); ii) shedding light on how external drivers (e.g. global changes) affect community dynamics in natural habitats (Bernhardt Römermann et al. 2015; Newbold et al. 2015) and iii) assessing relationships between community stability over time and the delivery of ecosystem services (Isbell et al. 2018). Reliable answers to these questions can only be provided by drawing upon ecological data collected at several points in time using consistent sampling procedures. For plant communities, long-term data collected by regularly sampling permanent plots probably constitute the most precise approach to detect temporal changes at the local scale (Bakker et al. 1996; Damgaard 2019; de Bello et al. 2020). First, due to their geographical position being kept “fixed” in the field, permanent plots prevent relocation bias, i.e. the error derived from trying to find the original plot location. This bias is inherent in vegetation resurveys (Verheyen et al. 2018). Second, the repeated collection of vegetation data from permanent plots provides broad benefits to our understanding of vegetation change, including the means to track detailed successional trajectories, monitor species interactions over time and assess the stability of the community as a whole. For this reason, permanent plots have been listed among the six most important developments in vegetation science over the past three decades (Chytrý et al. 2019).

In recent decades, vegetation science has benefited from the development and maintenance of large vegetation databases (Dengler et al. 2011). Historical vegetation relevés performed by early vegetation ecologists, together with recent vegetation plot data stemming from regional, but also national or continental research and survey projects, have been carefully assembled and digitally archived in the context of centralized initiatives (Chytrý et al. 2016; Wiser 2016; Bruelheide et al. 2019; Sabatini et al. 2021). Such global collections of vegetation plot data are essential to investigate macroecological patterns and provide spatially meaningful answers to global issues, i.e. to effectively perform global scale biodiversity research. In this context, a comparable effort specifically aimed at assembling and maintaining global databases built on time-series of vegetation data is urgently needed to lay a common ground for future studies focusing on i) providing global estimates of changes in plant diversity trends over time; ii) monitoring the conservation status of natural habitats over time or iii) assessing the stability of ecosystem functions and services. To the best of our knowledge, the BioTIME initiative (Dornelas et al. 2018) represents the most important global collection of biodiversity time-series including abundance records measured in species assemblages belonging to the marine, freshwater and terrestrial environments. Yet, the powerful spatial representation of BioTIME has considerable limitations that include: i) an often limited length and/or periodicity of the time-series, which particularly affects vegetation data (29 datasets with at least 6 data points; ii) a poor focus on vegetation and terrestrial plant biodiversity (96 datasets, corresponding to about 27% of the whole database). Given that a high number of ecosystem functions and services strongly depend on plants (Maestre et al. 2012; van der Plas 2019), we deem it crucial for the fields of vegetation science and ecology to be able to rely on a consistent and standardized collection of datasets including high-quality time-series measured at regular intervals and specific to plant communities.

Based on these premises, we hereby present the LOng-Term Vegetation Sampling (LOTVS) initiative, a growing global collection of vegetation time-series derived from the regular (mostly, annual) monitoring of vascular plants in permanent plots. By promoting the use, and supporting the visibility of high-quality temporal data collected using permanent plots, LOTVS ultimately aims to provide the tools to ask relevant ecological questions across a number of taxa, ecosystems and regions. The LOTVS collection provides a platform for aggregating the currently disconnected datasets sampled around the world based on permanent plots. As such, researchers are welcome to contribute to and, based on a scientific proposal (see section 3.2.), use the available collection of data.

## 2. Description of LOTVS

As of April 2021, LOTVS encompasses 79 datasets (Fig. 1) for a total of 7789 vegetation time-series, collected using permanent plots that were monitored for a minimum of six, and a maximum of 99 years (first quantile: 10; mean: 17.5; median: 16; third quantile: 23 years; see Fig 2). The vast majority of LOTVS time-series has a fine-grained temporal resolution: measurements in permanent plots were taken on 10% to 100% of the temporal interval (with 100% meaning that plots were sampled at least annually; first quantile: 82.8%; mean: 87.4%; median: 100%; third quantile: 100%). A description of the single datasets can be found in Valencia et al. (2020; Supplementary Material). At present, LOTVS includes vegetation time-series specifically focused on herbaceous species and shrubs mostly belonging to grassland habitats, followed by mixed vegetation types (i.e. savannas and heathlands), shrublands, forest understoreys and other habitats (mostly salt marshes; Fig. 4A; see Box 1). Forest plots monitoring changes in long-term changes in tree species are, at the moment, excluded from LOTVS (but see section Perspectives). The LOTVS collection contains data from five continents (Fig. 1; but see here for an interactive map), although Europe and North America are so far the most represented areas. While Europe is the leading continent in terms of datasets (38 out of 79), North America hosts the majority of plots (almost 70%, distributed across 30 datasets).

**Box 1.**
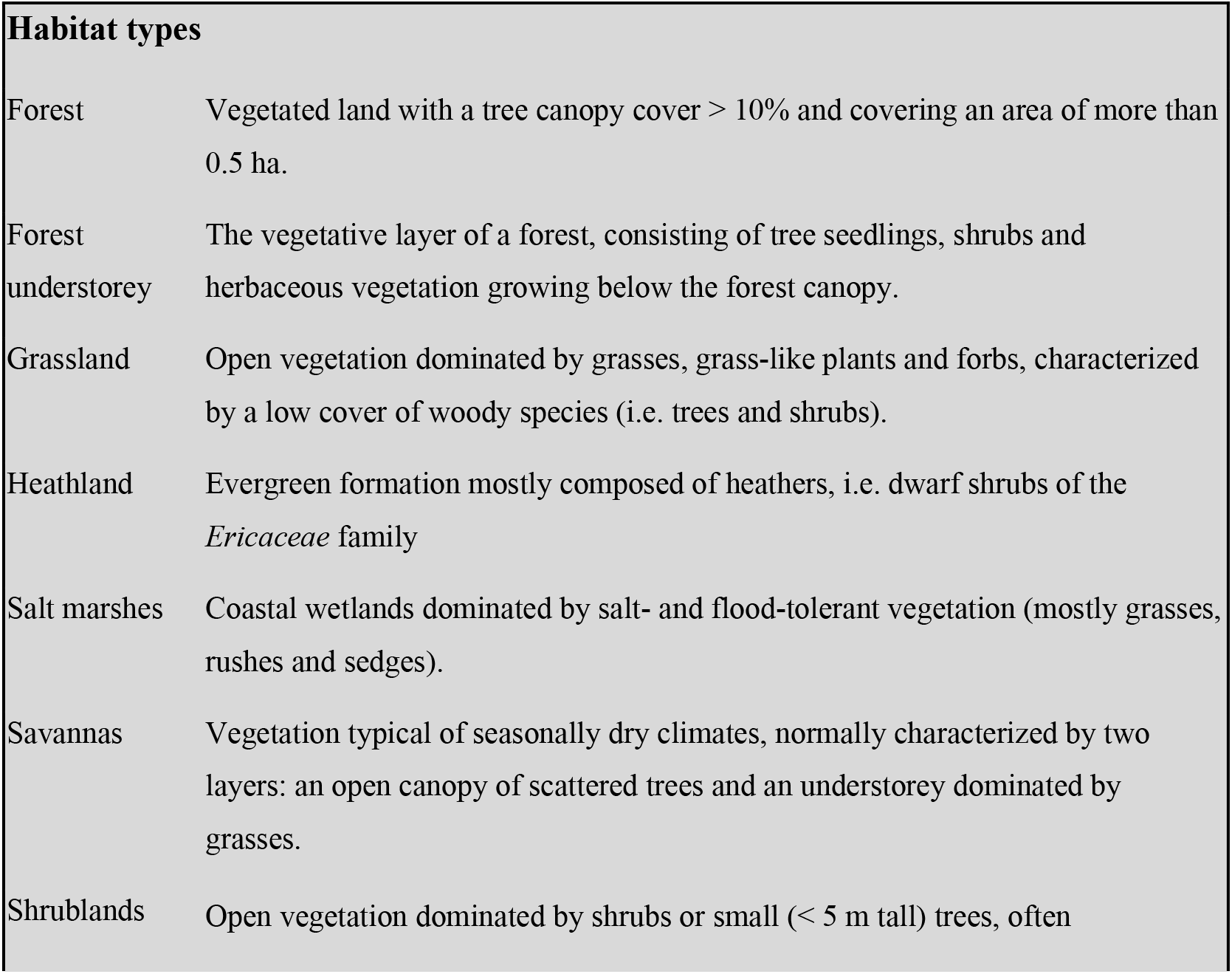
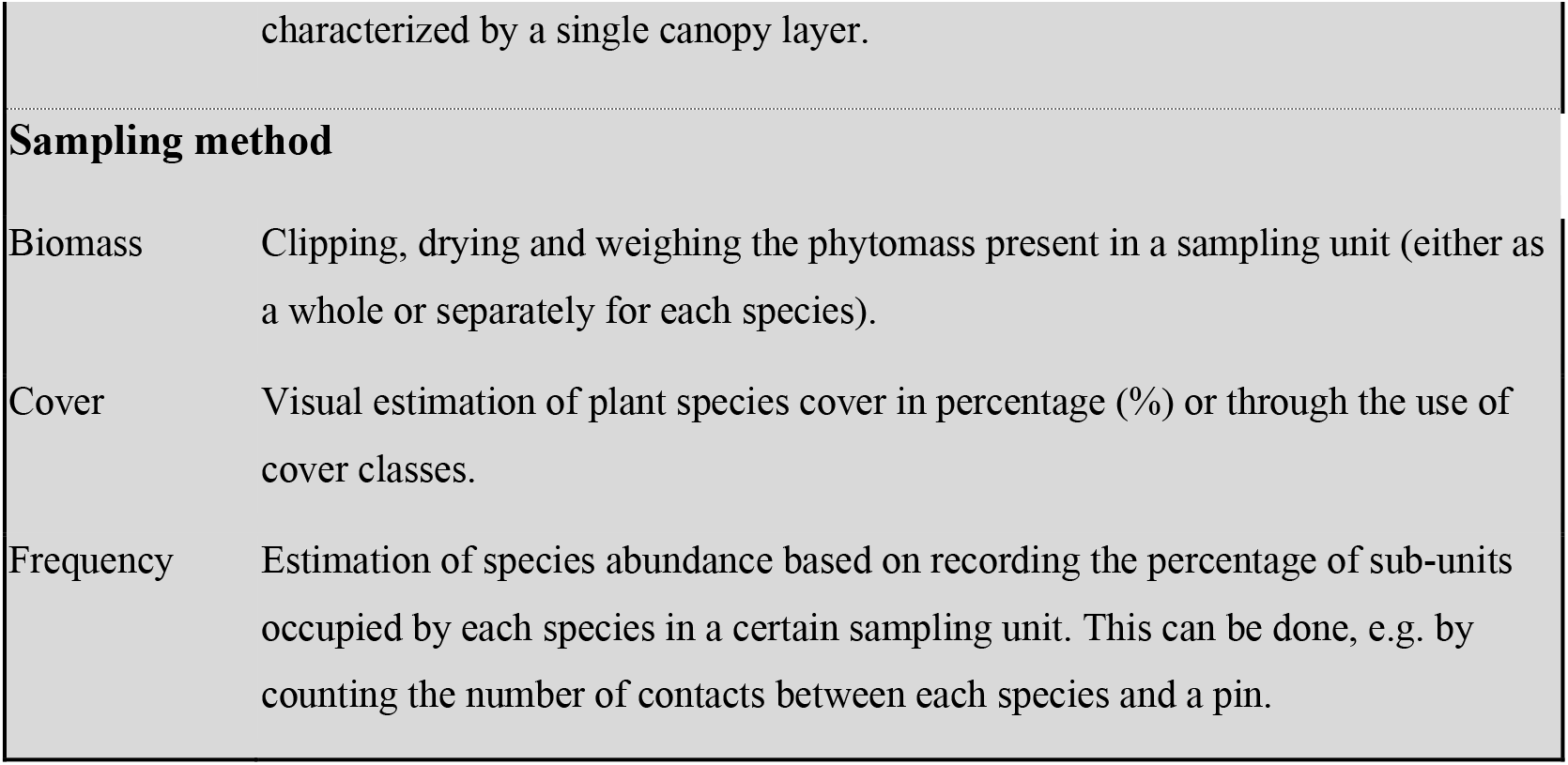
Working definitions of the main habitat types and sampling methods mentioned. Habitat definitions were adapted from the Convention of Biological Diversity website (https://www.cbd.int/forest/definitions.shtml and https://www.cbd.int/drylands/definitions), and from Goldstein & DellaSala (2020).

**Fig. 1.**
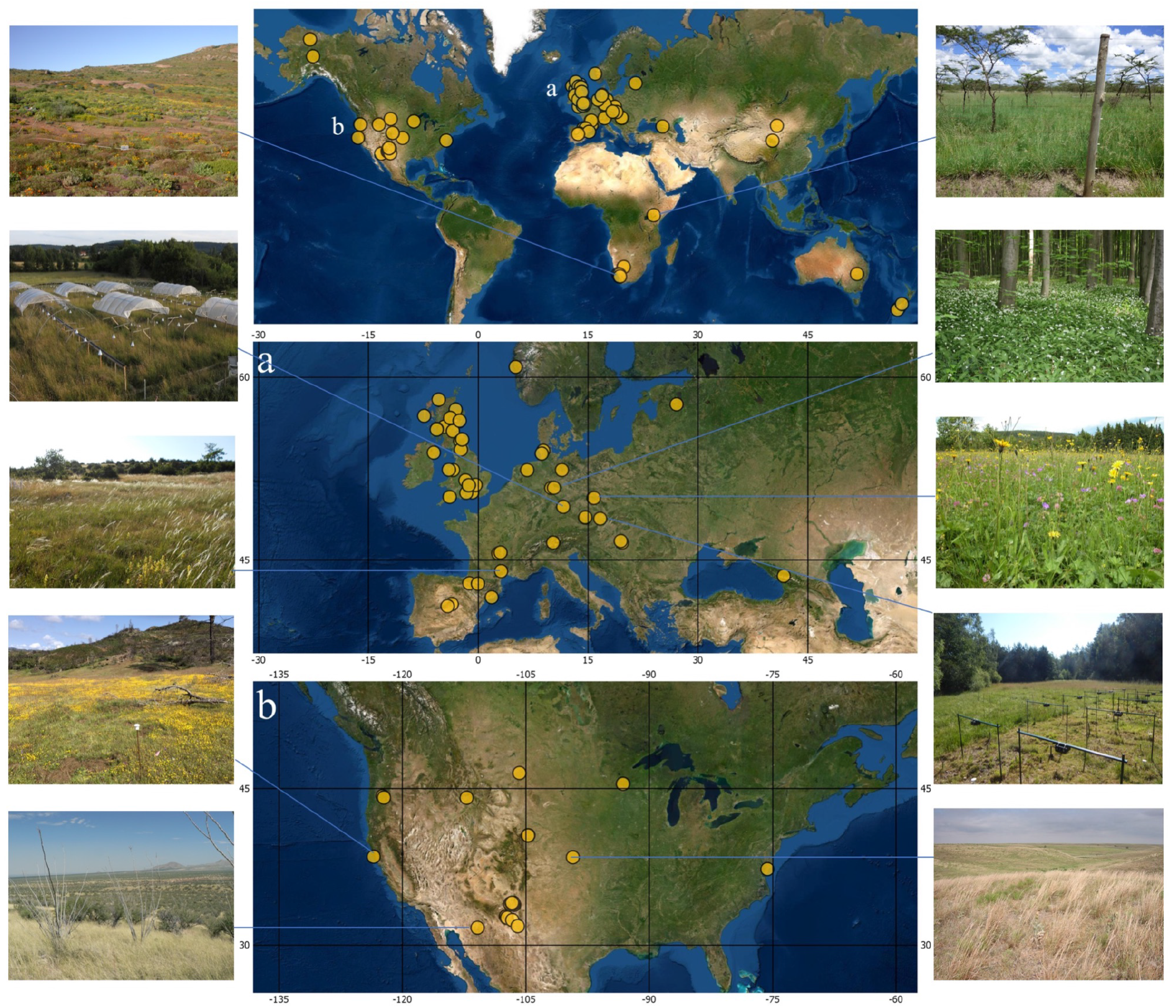
Map showing the geographical location of the 79 datasets included in LOTVS. A more detailed view is given for areas featuring a high density of datasets: a) Europe; b) North America. ESRI World Satellite Imagery was used as the base map. For a subset of sites, representative vegetation types are shown. Photos were taken at: (left, starting from the top): Soebatsfontein (South Africa), Bayreuth (Germany), Roquefort-sur-Soulzon (France), California (USA), Arizona (USA); (right, starting from the top): Laikipia (Kenya), Göttinger Wald (Germany), Krkonose Mountains (Czech Republic), Ohrazeni (Czech Republic), Kansas (USA).

**Fig. 2.**
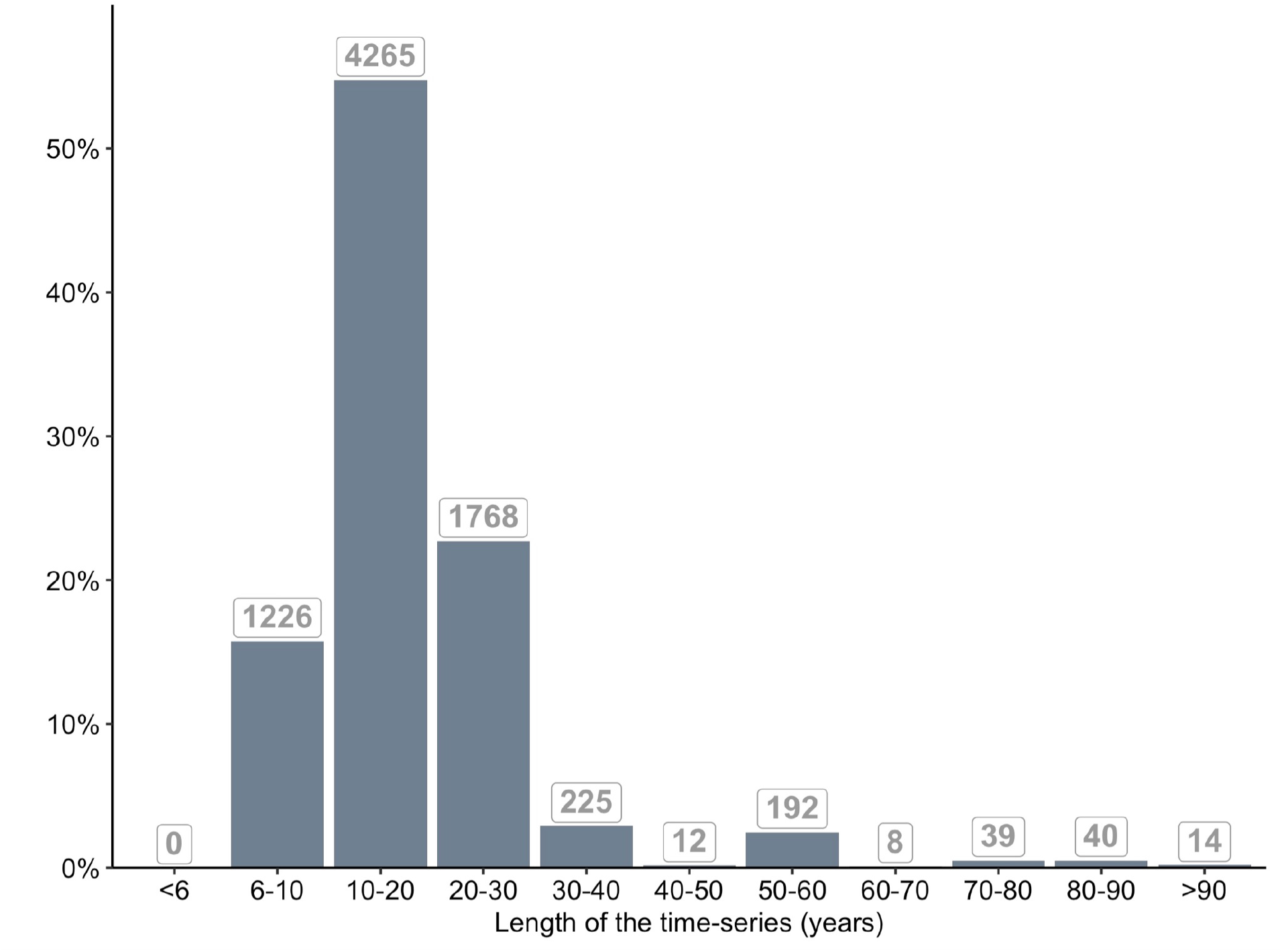
Distribution of the duration (in years) of the time-series included in LOTVS. The number of time-series within each class is reported above the bars.

Datasets included in LOTVS span a wide climatic gradient, their mean annual temperature ranging between −11.5 and 20.1°C, and their mean annual precipitation between 14 and 259.2 cm (source: WordClim 2; Fick & Hijmans 2017). As such, they are mostly included in the temperate seasonal forest, temperate grassland/desert and in the woodland/shrubland biomes (sensu Whittaker 1975; see Fig 3).

**Fig. 3.**
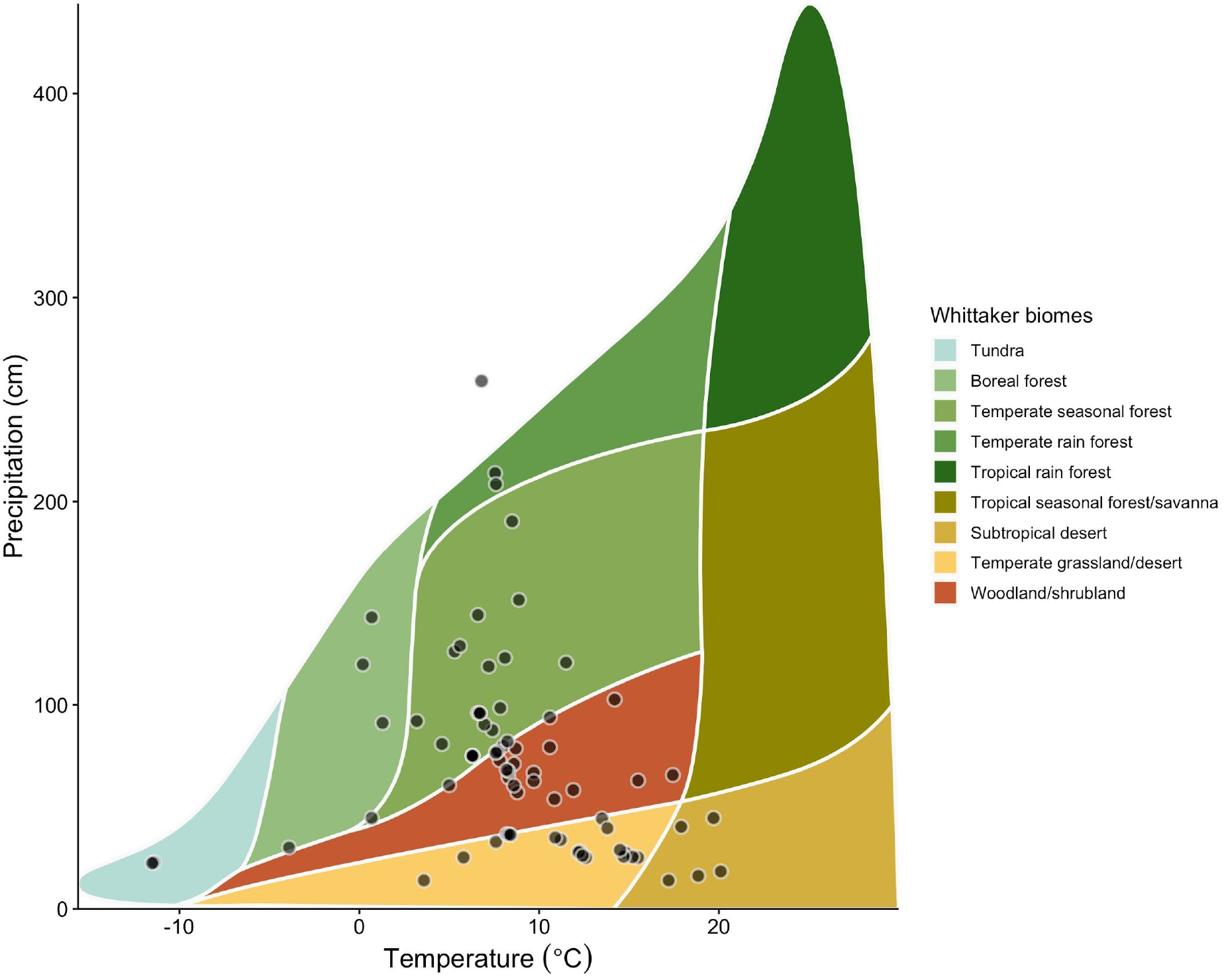
Climatic summary of the 79 datasets included in LOTVS. Mean annual temperature and mean annual precipitation are plotted on the x and y axes, respectively. Each dot represents mean climatic conditions characterizing each dataset. Dots are superimposed on Whittaker biomes (i.e. indicating potential vegetation; Whittaker, 1975), as redrawn from Ricklefs (2008). The plot was created using the R package “plotbiomes” (Stefan and Levin 2021).

**Fig. 4.**
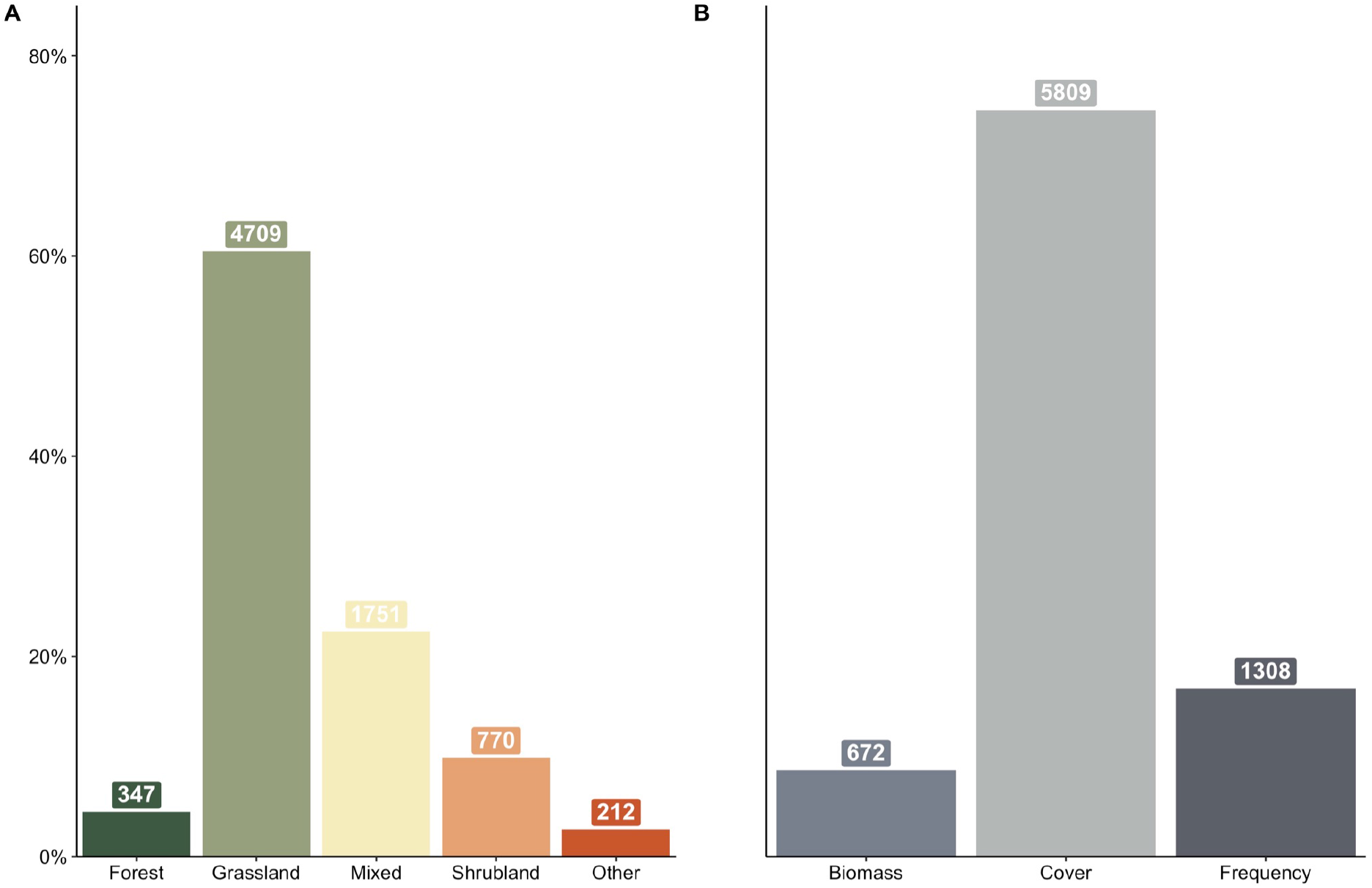
Habitat types (A) and sampling methods/approaches (B) covered in LOTVS. The number of time-series within each class is reported on top of the bars. See Box 1 for working definitions of habitat type and sampling methods.

In almost half of the permanent plots included in LOTVS (48.5%), vegetation has been subjected to experimental treatments manipulating abiotic or biotic conditions. The most frequent treatment types are herbivore exclusion, fertilization and grazing intensification (applied to ∼35, 24 and 17% of the treated plots, respectively; Fig. 5). Yet, even in the absence of such treatments, LOTVS includes plots subjected to management regimes, such as mowing or grazing, that are necessary to maintain traditional land-use in given habitats. It should be noted that in 30 datasets (corresponding to about 50% of the datasets within LOTVS) traditional management was maintained as a control comparison to novel treatments.

**Fig. 5.**
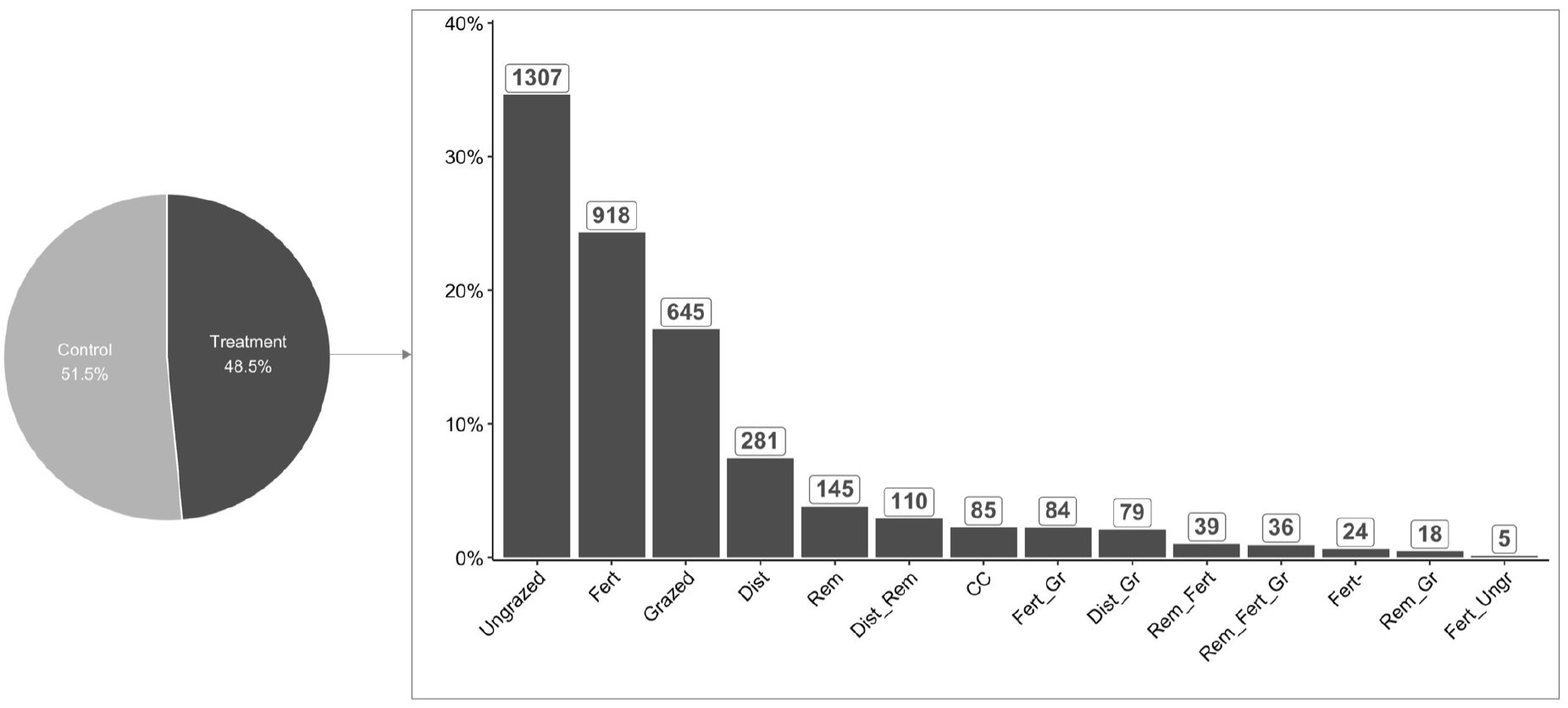
Left: pie chart describing the distribution of permanent plots included in the LOTVS collection according to the absence (“control”) or presence (“treatment”) of treatment. Right: distribution of the type of treatments present in LOTVS. The number of time-series within each treatment is reported on top of the bars. Names for the treatments were abbreviated as follows: “Ungrazed”: grazing exclosure; “Fert”: fertilization; “Grazed”: grazing; “Dist”: disturbance; “Rem”: removal; “Dist_Rem”: disturbance + removal; “CC”: climate change; “Fert_Gr”: fertilization + grazing; “Dist_Gr”: disturbance + grazing; “Rem_Fert”: removal + fertilization; “Rem_Fert_Gr”: removal + fertilization + grazing; “Fert-”: decrease of productivity; “Rem_Gr”: removal + grazing; “Fert_Ungr”: fertilization + grazing exclosures.

Permanent plots in the LOTVS collection are surveyed using different techniques. The vast majority (∼85%) are quadrat plots, but line transects and quadrat plots arranged along a transect are also present. Plot size ranges from 0.04 to 400 m^2^; ∼ 80% of the plots range from 0.04 to 1.25 m^2^, with 1 m^2^ being the most frequent (49%) plot size in the collection. Information on plot size is missing for 90 plots, corresponding to 0.8% of the whole LOTVS collection. The method used to quantify species abundance also varies among the 79 datasets (Fig. 4B). The most frequent approach uses visual estimation of species cover (75%) followed by recording the frequency of individual species across a given number of subplots (17%) and third by the collection of aboveground biomass for each species in the plot (9%). In most cases, biomass clipping is intended to mimic mowing. All plots included in LOTVS were permanently marked in the field; geographic coordinates are available for either specific plots or for unique localities of each dataset. Besides estimating plant species abundance, some of the datasets within LOTVS originally included information about bare ground cover or the abundance of other taxa (e.g. bryophytes, lichens). However, in an effort to provide a standardized and consistent tool, this information was removed from individual datasets so that recorded taxa within LOTVS now only include vascular plants.

Taxonomic standardization is fundamental when compiling vegetation databases. On the one hand, it is a necessary step to addressing the issue of nomenclature redundancy caused by the multitude of synonyms characterizing botanical literature (Kalwij 2012); on the other hand, it allows a later integration of data with ancillary information linked to taxonomic entities, e.g. species functional traits. Original datasets included in LOTVS show considerable variation in the chosen taxonomical references reflecting different regional and national traditions as well as the time when the work was undertaken. As such, a taxonomic standardization was deemed necessary. To do this, we standardized the nomenclature of plant species following The Plant List, currently the most widely used global reference list (Kalwij 2012). This was done in R (R Core Team 2019) using the package “Taxonstand” (Cayuela et al. 2019), which allows the automatic standardization of plant names by running an internal query to The Plant List (http://www.theplantlist.org) and returning standardized species names, eventually resolving synonyms and homogenizing intraspecific taxonomic entities to the level of species.

## 3. Data usage

As an unprecedented collection of vegetation time-series, LOTVS has a huge potential to support groundbreaking research in the fields of vegetation science, plant ecology and temporal ecology. It should be noted, though, that installing and maintaining permanent plots is a very time- and resource-consuming task, and thus a powerful collection of vegetation plots such as LOTVS can only arise from collaborative efforts that stem from an impressive amount of work carried out by many data contributors, whose effort must be acknowledged. To this end, anyone can contribute to LOTVS with original data, as long as the data fully comply with LOTVS requirements (see section 3.2.). At the same time, data included in LOTVS can be requested and used following a simple procedure that is intended to maximize transparency and support well-grounded research projects.

### 3.1. Contributing data

Detailed information on how to contribute to LOTVS, as well as specific data requirements can be found on the dedicated website https://lotvs.csic.es/contribute/. LOTVS welcomes datasets including vegetation time-series collected from permanent plots with a fixed (i.e. permanently marked) geographical position in the field, possibly replicated in space, maintained for a minimum of 6 years and sampled at regular (preferably annual) intervals. In principle, the time-series should be continuous, i.e. gaps should not be present. However, exceptions are allowed, provided that observations for some years are only missing for a reduced number of plots. In cases of missing years for all of the permanent plots, the new dataset can only be incorporated to LOTVS if a) only a very limited number of years is missing and b) their distribution within the time-series is irregular, i.e. LOTVS is not intended to accept permanent plots that, according to their original scope, are only sampled every *n* years. Following these requirements, we are also not looking to include data collected in the context of so-called resurveying studies, where historic vegetation plots are revisited after a longer time period and re-recorded. In fact, these are the subject of other databases (e.g. the ReSurveyEurope initiative, http://euroveg.org/eva-database-re-survey-europe). Also, whereas data collected using different sampling approaches are welcomed (e.g. visual estimation of species cover, biomass, frequency, number of individuals), the sampling approach should be as consistent as possible over time. LOTVS does not plan, at present, to incorporate permanent plots that only record species occurrence (i.e. presence/absence data). To be included in LOTVS, permanent plots should be preferably representative of natural or semi-natural vegetation. The former can be defined as vegetation that developed in absence of human influence and/or has long been left undisturbed by humans; as to the latter, its existence and maintenance depend on human practices (e.g. grazing, mowing) carried out for either production, conservation, or a mix of the two purposes. As such, time-series data recorded from artificial seed mixtures such as those sown in biodiversity experiments are not currently accepted.

### 3.2. Requesting data

Because of the effort needed to maintain permanent plots in time and collect temporal vegetation data (Mills et al. 2015), access to individual time-series or datasets included in LOTVS is governed by a data policy that allows data owners and contributors to remain in full control of their data if they so choose (see https://lotvs.csic.es/contribute/ for detailed information). The availability and access of single datasets depend on the choice of individual data owners and contributors, who decide the accessibility level of their data (from “restricted data”, i.e. data are only usable upon consent from data owners/contributors, that should be expressed each time their dataset is requested; to “free data” i.e. data that are freely available to use through the LOTVS platform). We note that about 40% of LOTVS datasets are publicly available, either because they belong to Long-Term Ecological Research (LTER) Programs, or because they were archived and published by their data owners. Also, several of the LOTVS datasets are publicly available in their own right, via contact with their owners. Depending on the accessibility level specified, data owners and contributors hold (or not, in case of freely usable data) the right to request authorship on eventual publications based on the proposal submitted by the applicants when they request the data. In all cases, to request data included in LOTVS, a short and sound scientific proposal describing the aims of the project and the type of data required should be prepared and submitted to the LOTVS’ Supervising Committee. This process is intended to i) minimize conceptual overlap of proposals addressing highly similar research questions and ii) make sure that all data owners are informed about the possible use of their data and are free to decline it if they wish so.

#### Perspectives

In its present form, LOTVS includes vegetation time-series for almost 8000 permanent plots installed and maintained in natural and semi-natural plant communities worldwide. Still, as we explained in section 2, the geographical representation of both individual datasets and permanent plots is not homogeneous; it is biased towards Europe and North America, and many habitats, such as forest habitats, are strongly underrepresented. In order to promote a more equal representation in terms of geographical areas and habitats, one of the goals of this paper is to encourage new datasets including time-series recorded using permanent plots located in currently underrepresented continents such as Africa, Asia, Australia and South America. Similarly, time-series recorded in forests, tundra and coastal areas would be particularly welcome. Concerning forests, it should be noted that in its current version LOTVS only includes permanent plots located in the understory layer, while we look forward to increasing the representation of actual forest time-series. Furthermore, we are very interested in datasets featuring spatial as well as temporal replication (minimum 6 years as mentioned above) to disentangle the differences between temporal and spatial changes. Finally, to broaden the range of potential applications of LOTVS (and in line with what has been done by other global initiatives, see Bruelheide et al. 2019), we are planning to integrate it with information on environmental variables (climate, micro-climate) and species functional traits. Such integration will eventually allow users direct access to ancillary data crucial to explore the evolution of different facets of diversity over time (Monnet et al. 2014; Sperandii et al. 2021).

## Conclusions

LOTVS possibly represents the largest collection of temporally fine-grained vegetation time-series made accessible to the research community derived from permanent plots addressing the study of plant communities through time. As such, LOTVS can be highly useful to perform timely research on a wide range of topics in the field of vegetation science: investigating patterns and drivers of ecological succession in natural plant communities, quantifying vegetation changes through time, as well as assessing community stability and identifying its mechanisms. At the same time, because it includes a considerable proportion of permanent plots subjected to some kind of treatment (e.g. grazing, fertilization etc.), LOTVS can also support the development of large-scale studies aiming to understand how temporal dynamics are affected by different treatments in the context of global changes. Last, but not least, we believe LOTVS could also serve as a valuable resource to conduct methodological research addressing topics related to, for example, methods to quantify dissimilarity through time and their partitioning (Baselga 2010; Legendre & Condit 2015) or quantitative approaches to investigate community dynamics and more specifically, stability.

## References

Bakker, J. P., Olff, H., Willems, J. H., & Zobel, M. 1996. Why do we need permanent plots in the study of long term vegetation dynamics? Journal of Vegetation Science 7(2): 147–156.

Baselga, A. 2010. Partitioning the turnover and nestedness components of beta diversity. Global Ecology and Biogeography 19(1): 134–143.

Bernhardt Römermann, M., Baeten, L., Craven, D., De Frenne, P., Hédl, R., Lenoir, J., … & Verheyen, K. 2015. Drivers of temporal changes in temperate forest plant diversity vary across spatial scales. Global Change Biology 21(10): 3726–3737.

Blowes, S. A., Supp, S. R., Antão, L. H., Bates, A., Bruelheide, H., Chase, J. M., … & Dornelas, M. 2019. The geography of biodiversity change in marine and terrestrial assemblages. Science 366(6463): 339–345.

Bradshaw, C. J., Ehrlich, P. R., Beattie, A., Ceballos, G., Crist, E., Diamond, J., … & Blumstein, D. T. 2021. Underestimating the challenges of avoiding a ghastly future. Frontiers in Conservation Science: 1–9.

Brondizio E.S., Settele J., Díaz S., & Ngo H.T.(Eds.). 2019. IPBES: Global assessment report on biodiversity and ecosystem services of the Intergovernmental Science Policy Platform on Biodiversity and Ecosystem Services. Bonn, Germany: IPBES Secretariat.

Bruelheide, H., Dengler, J., Jiménez Alfaro, B., Purschke, O., Hennekens, S. M., Chytrý, M., … & Tang, Z. 2019. sPlot–A new tool for global vegetation analyses. Journal of Vegetation Science 30(2): 161–186.

Cayuela, L., Macarro, I., Stein, A. and Oksanen, J. 2019. Taxonstand: Taxonomic Standardization of Plant Species Names. R package version 2.2. https://CRAN.R-project.org/package=Taxonstand.

Ceballos G., Ehrlich P.R., Barnosky A.D., García A., Pringle R.M., & Palmer T.M. 2015. Accelerated modern human–induced species losses: Entering the sixth mass extinction. Science Advances 1(5): e1400253.

Chytrý M., Hennekens S.M., Jiménez Alfaro B., Knollová I., Dengler J., Jansen F., … & Yamalov S. 2016. European Vegetation Archive (EVA): an integrated database of European vegetation plots. Applied Vegetation Science 19(1): 173–180.

Chytrý M., Chiarucci A., Pärtel M. and Pillar V.D. 2019. Progress in vegetation science: trends over the past three decades and new horizons. Journal of Vegetation Science 30: 1–4.

Convention on Biological Diversity. 2020. Global Biodiversity Outlook. Montréal, QC: Secretariat of the Convention on Biological Diversity.

Damgaard C. 2019. A critique of the space-for-time substitution practice in community ecology. Trends in Ecology & Evolution 34: 416–421.

de Bello F., Valencia E., Ward D., & Hallett L. 2020. Why we still need permanent plots for vegetation science. Journal of Vegetation Science 31(5): 679–685.

Dengler J., Jansen F., Glöckler F., Peet R.K., De Cáceres M., Chytrý M., … & Spencer N. 2011. The Global Index of Vegetation Plot Databases (GIVD): a new resource for vegetation science. Journal of Vegetation Science 22(4): 582–597.

Dornelas M., Antao L.H., Moyes F., Bates A.E., Magurran A.E., Adam D., … & Murphy G. 2018. BioTIME: A database of biodiversity time series for the Anthropocene. Global Ecology and Biogeography 27(7): 760–786.

Fick S.E., & Hijmans R.J. 2017. WorldClim 2: new 1 km spatial resolution climate surfaces for global land areas. International Journal of Climatology, 37(12): 4302–4315.

Goldstein, M.I., & DellaSala, D.A. 2020. Encyclopedia of the World’s Biomes. Elsevier.

Gonzalez A., Cardinale B.J., Allington G.R., Byrnes J., Arthur Endsley K., Brown D.G., … & Loreau M. 2016. Estimating local biodiversity change: a critique of papers claiming no net loss of local diversity. Ecology 97(8): 1949–1960.

Hillebrand H., Blasius B., Borer E.T., Chase J.M., Downing J.A., Eriksson B.K., … & Ryabov A. B. 2018. Biodiversity change is uncoupled from species richness trends: Consequences for conservation and monitoring. Journal of Applied Ecology 55(1): 169–184.

Jones S.K., Ripplinger J., & Collins S.L. 2017. Species reordering, not changes in richness, drives long term dynamics in grassland communities. Ecology Letters 20(12): 1556–1565.

Kalwij, J.M. (2012). Review of ‘The Plant List, a working list of all plant species’. Journal of Vegetation Science 23(5): 998–1002.

Legendre P., & Condit R. 2019. Spatial and temporal analysis of beta diversity in the Barro Colorado Island forest dynamics plot, Panama. Forest Ecosystems 6(1): 1–11.

Maestre F.T., Quero J.L., Gotelli N.J., Escudero A., Ochoa V., Delgado-Baquerizo M., … & Zaady E. 2012. Plant species richness and ecosystem multifunctionality in global drylands. Science 335(6065): 214–218.

Mills J.A., Teplitsky C., Arroyo B., Charmantier A., Becker P.H., Birkhead T.R., … & Zedrosser A. 2015. Archiving primary data: solutions for long-term studies. Trends in Ecology & Evolution 30(10): 581–589.

Monnet A.C., Jiguet F., Meynard C.N., Mouillot D., Mouquet N., Thuiller W., & Devictor V. 2014. Asynchrony of taxonomic, functional and phylogenetic diversity in birds. Global Ecology and Biogeography 23(7): 780–788.

Newbold T., Hudson L.N., Hill S.L., Contu S., Lysenko I., Senior R.A., … & Purvis A. 2015. Global effects of land use on local terrestrial biodiversity. Nature 520(7545): 45–50.

Pereira H.M., Navarro L.M., & Martins I.S. 2012. Global biodiversity change: the bad, the good, and the unknown. Annual Review of Environment and Resources 37: 25–50.

Pimm S.L., Jenkins C.N., Abell R., Brooks T. M., Gittleman J.L., Joppa L.N., … Sexton J.O. 2014. The biodiversity of species and their rates of extinction, distribution, and protection. Science 344: 987.

R Core Team. 2019. R: A language and environment for statistical computing. R Foundation for Statistical Computing, Vienna, Austria. URL https://www.R-project.org/.

Ricklefs R.E. 2008. The economy of nature. Macmillan.

Sabatini F.M., Lenoir J., Hattab T., Arnst E., Chytrý M., Dengler J., … Bruelheide H. 2021. sPlotOpen – An environmentally-balanced, open-access, global dataset of vegetation plots. Global Ecology and Biogeography. https://doi.org/10.1111/geb.13346.

Sperandii M.G., Barták V., Carboni M., & Acosta A.T.R. 2021. Getting the measure of the biodiversity crisis in Mediterranean coastal habitats. Journal of Ecology 109(3): 1224–1235.

Stefan V. and Levin S. 2021. plotbiomes: Plot Whittaker biomes with ggplot2. R package version 0.0.0.9001.

Valencia E., de Bello F., Galland T., Adler P.B., Lepš J., E-Vojtkó A.., … & Götzenberger L. 2020. Synchrony matters more than species richness in plant community stability at a global scale. Proceedings of the National Academy of Sciences 117(39): 24345–24351.

van der Plas F. 2019. Biodiversity and ecosystem functioning in naturally assembled communities. Biological Reviews 94(4): 1220–1245.

Verheyen K., Bažány M., Chećko E., Chudomelová M., Closset Kopp D., Czortek P., … & Baeten, L. 2018. Observer and relocation errors matter in resurveys of historical vegetation plots. Journal of Vegetation Science 29(5): 812–823.

Whittaker R.H. 1975. Communities and Ecosystems, 2nd ed., Macmillan, New York.

Wiser S.K. 2016. Achievements and challenges in the integration, reuse and synthesis of vegetation plot data. Journal of Vegetation Science 27(5): 868–879.

